# Deficient synaptic neurotransmission results in a persistent sleep-like cortical activity across vigilance states in mice

**DOI:** 10.1101/2023.05.11.540034

**Authors:** Mathilde C. C. Guillaumin, Christian D. Harding, Lukas B. Krone, Tomoko Yamagata, Martin C. Kahn, Cristina Blanco-Duque, Gareth T. Banks, Patrick M. Nolan, Stuart N. Peirson, Vladyslav V. Vyazovskiy

## Abstract

Growing evidence suggests that brain activity during sleep, as well as sleep regulation, are tightly linked with synaptic function and network excitability at the local and global levels. We previously reported that a mutation in synaptobrevin 2 (*Vamp2*) in restless (*rlss*) mice results in a marked increase of wakefulness and suppression of sleep, in particular REM sleep (REMS) as well as increased consolidation of sleep and wakefulness. In the current study, using finer-scale *in vivo* electrophysiology recordings, we report that spontaneous cortical activity in *rlss* mice during NREM sleep (NREMS) is characterised by an occurrence of abnormally prolonged periods of complete neuronal silence (OFF-periods), often lasting several seconds, similar to the burst suppression pattern typically seen under deep anaesthesia. Increased incidence of prolonged network OFF-periods was not specific to NREMS, but also present in REMS and wake in *rlss* mice. Slow-wave activity (SWA) was generally increased in *rlss* mice, while higher frequencies including theta-frequency activity were decreased, further resulting in diminished differences between vigilance states. The relative increase in SWA after sleep deprivation was attenuated in *rlss* mice, suggesting either that *rlss* mice experience persistently elevated sleep pressure, or, alternatively, that the intrusion of sleep-like patterns of activity into awake state diminishes the accumulation of sleep drive. We propose that deficit in global synaptic neurotransmitter release leads to ‘state inertia’, reflected in an abnormal propensity of brain networks to enter and remain in a persistent ‘default state’ resembling coma or deep anaesthesia.

## INTRODUCTION

Despite recent advances in our understanding of neurophysiological and molecular mechanisms of sleep-wake control, it remains unclear how sleep and wake alternation is regulated at a brain-wide scale, and whether such fundamental aspects of neuronal function as synaptic release have a role to play in that respect.

The restless (*rlss*) mouse line, obtained via a forward genetics sleep screen and possessing a mutation in synaptobrevin 2/*Vamp2* (a gene encoding the key synaptic-release machinery protein VAMP2) showed an array of sleep and behavioural phenotypes (Banks et al., 2020), including a decreased amount of NREM sleep (NREMS) in the dark phase, alongside an overall decrease in REM sleep (REMS) time. More strikingly, the number of sleep and wake episodes and of state transitions was markedly reduced in *rlss* mice, correlating with increased duration of episodes in all vigilance states compared to littermate controls. The probability to transition from one state to the next in normal conditions was found to be markedly reduced in *rlss* mice, leading to a general ‘vigilance state inertia’.

The mutation in *Vamp2* results in a markedly decreased neurotransmitter release probability in *rlss* mice, altering synaptic transmission (Banks et al., 2020). These findings raised two questions. First, why did this state-transitioning probability decrease for all states, and in both directions between two given states (except NREMS-to-wake transitions)? Secondly, what are the underpinning mechanisms of the decrease in time spent in REMS? Having shown changes in the architecture of vigilance states (Banks et al., 2020), we now take a different approach and look at the ‘quality’ of the states themselves, to address the question of whether changes in state dynamics and architecture between *rlss* and wild-type (WT) mice induce qualitative differences between states.

EEG slow-wave activity (SWA), which corresponds to the EEG power in the slow-wave frequency range (0.5-4 Hz) in NREMS, has been a useful measure to evaluate sleep-wake dynamics. Slow-waves tend to dominate the EEG during NREMS and can be recorded across the entire cortical surface (Massimini et al., 2004; Steriade et al., 1993). Both animal and human studies have shown that SWA levels are closely linked to sleep-wake history, where SWA levels at the onset of NREMS positively correlate with preceding time spent awake, and levels subsequently decrease during NREMS (Achermann and Borbely, 2003; Borbely et al., 1984). SWA has therefore been traditionally used as a marker of sleep pressure (Borbely et al., 1981; Dijk et al., 1987). However, EEG records global signals across the cortical surface. Local field potentials (LFP) and multi-unit activity (MUA) instead allow us to look at sleep-wake dynamics at a more local level. In addition, as the initiation of slow-waves (depth positive/surface negative deflections) correspond to periods of neuronal silence (Calvet et al., 1973; Contreras and Steriade, 1995; Harding et al., 2023; Mukovski et al., 2007; Steriade et al., 1993; Steriade et al., 2001; Vyazovskiy et al., 2009), which also depend on the levels of network synchrony (Vyazovskiy et al., 2009), multi-unit activity allows us to investigate the underlying patterns of neuronal silence (‘OFF-periods’) (Harding et al., 2023).

In the present study, we used EEG, LFP and MUA recordings, combined with sleep deprivation and auditory stimulation paradigms, to gain a better understanding of the sleep depth of *rlss* mice. Auditory stimuli are commonly used to investigate sleep depth, sensory dissociation during sleep and arousal thresholds from NREMS and REMS (Hayat et al., 2022; Neckelmann and Ursin, 1993; Nir et al., 2015; Rechtschaffen et al., 1966; Wimmer et al., 2012). We investigated the spectral signature of global sleep and wake states, in particular at state transitions. We further explored the neuronal dynamics underlying the marked behavioural changes observed, and, using a mathematical model of sleep regulation (Guillaumin et al., 2018; Thomas et al., 2020), we examined the homeostatic regulation of sleep in *rlss* mice. We found that in *rlss* mice, the boundaries between electrophysiological signatures of vigilance states were less distinct than in control animals, and that sleep homeostasis dynamics were altered, with *rlss* mice being slower to dissipate accumulated sleep pressure. This was not associated, however, with an altered responsiveness to external sensory stimuli such as a sound.

This study suggests that impaired neurotransmitter release leads to an increased propensity to revolve around a ‘default state’ of cortical activity (Bandarabadi et al., 2020; Buzsáki, 2006; Lemieux et al., 2014; Sanchez-Vives et al., 2017; Sanchez-Vives and Mattia, 2014; Timofeev et al., 2020), with important consequences for global and local sleep architecture and sleep homeostasis.

## RESULTS

### *rlss* mice show increased propensity for hybrid states of vigilance

Using EEG combined with LFP recordings in freely-moving mice from the motor (M1) and visual (V1) cortical areas (**Figure 1a**), we found that *rlss* mice have an increase in spectral power in the very low frequencies in NREMS and REMS and a decrease in the higher frequency bins in REMS (EEG only), NREMS and Wake (**Figures 1b and S1a**). Interestingly, the differences between *rlss* and WT mice’ spectral signatures diverged between the frontal EEG and LFP signals in REMS, with an increase in the very-low delta band and theta (and higher) frequencies in the frontal REMS EEG, and an increase in the upper delta band in the frontal LFP in *rlss* mice compared to WT mice (**Figure 1b, middle**). Changes in the EEG occipital derivation paralleled the ones observed in the frontal EEG (**Figure S1a**); therefore, as LFP signals were recorded from M1, we also focused on EEG signals from this region (frontal). Local cortical dynamics thus revealed a disparity between local intracortical and brain surface (‘global’) signatures of REMS in *rlss* mice.

**Figure 1.**
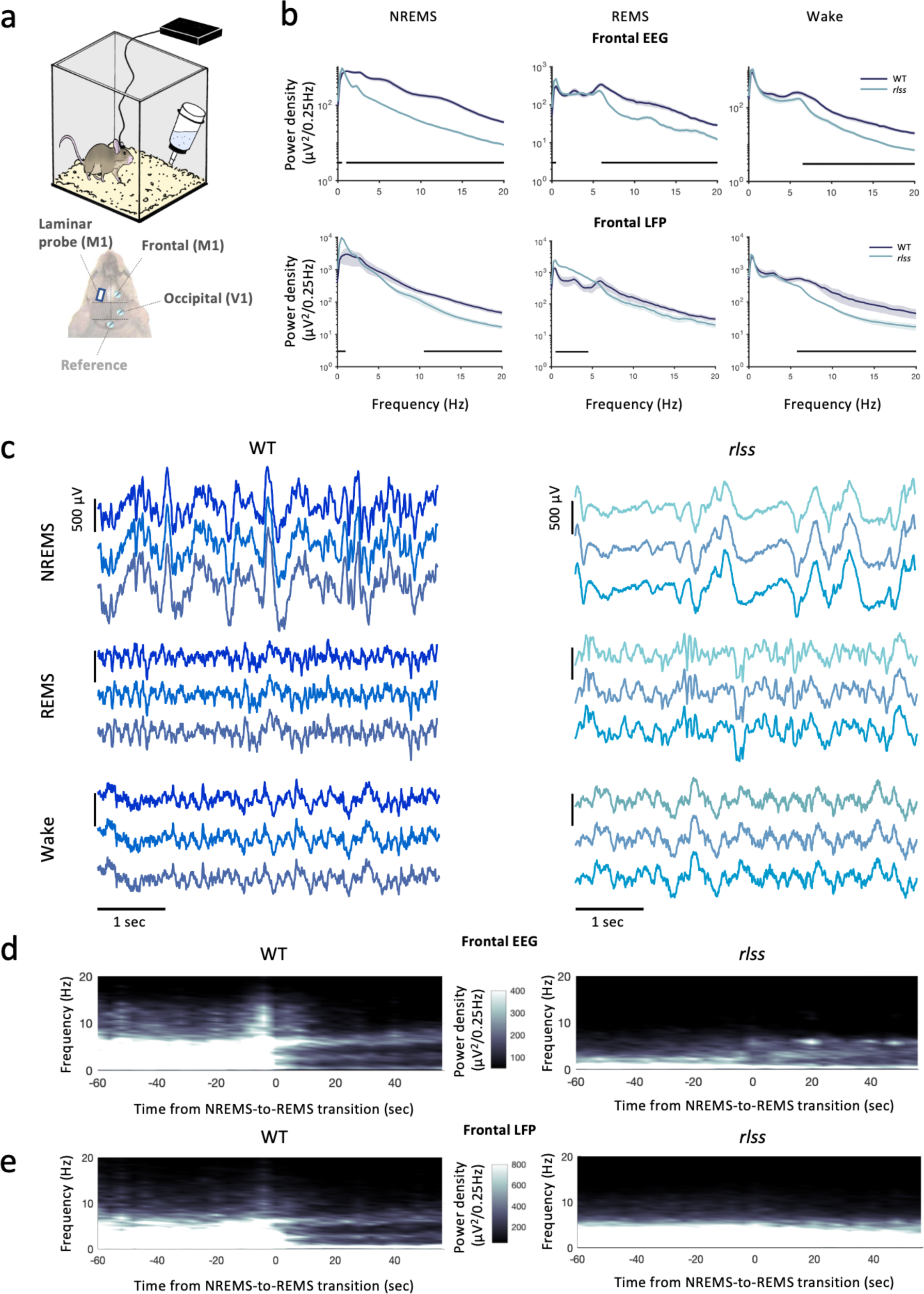
*rlss* mice show an increased level of similarity between vigilance state signatures. a) Schematic of recording set up. M1: primary motor cortex; V1: primary visual cortex. b) Frontal EEG (upper panels) and LFP (lower panels) power spectra during NREMS, REMS and wake in WT and *rlss* mice. Black bars at the bottom indicate the frequency bins for which the power is significantly different (p<0.05) between WT and *rlss* mice. Values shown as mean ± SEM. c) Examples of three LFP traces from a WT mouse (left) and *rlss* mouse (right) during NREMS, REMS and wake. d) Evolution of the power spectra in the frontal EEG at NREMS-to-REMS transitions averaged across WT (left) and *rlss* (right) mice. e) As in d), but for LFP signals. Note that here n(WT)=3. Unless otherwise stated, n(WT)=5 and n(*rlss*)=5.

LFP signals in *rlss* mice were also characterised by a decrease in their amplitude, especially during NREMS, when traces were reminiscent of the burst suppression patterns that are typically observed during general anaesthesia, but also coma, hypothermia and some early infantile pathologies (Brown et al., 2010; Clark and Rosner, 1973; Huang et al., 2021; Shanker et al., 2021) (**Figure 1c and Movie 1**). Such EEG/LFP signatures across vigilance states indicate that sleep inertia not only means fewer state transitions in *rlss* mice (Banks et al., 2020), but also that they display an intrusion of default-mode-like traits (Sanchez-Vives et al., 2017) across all states of vigilance, with the well-known spectral signatures of NREMS, REMS and wake being affected. The boundaries between states’ typical signatures appear less distinct in *rlss* mice, due to an increased power in the very slow frequencies not only in NREMS, but also in REMS and wake, associated with a decreased theta power in REMS in *rlss* mice. At a finer time resolution, looking at spectrograms at the transitions from NREMS to REMS, we found typical differences between *rlss* and WT mice. Blurred boundaries between states in *rlss* mice appeared all the more evident, with the drop in spectral power in the slow-frequency range (0.5-4 Hz) that is usually observed at NREMS-to-REMS transitions being significantly reduced in *rlss* mice (**Figures 1d,e**).

### *rlss* mice show prolonged periods of neuronal silence across all vigilance states, and especially during NREMS, reminiscent of burst suppression patterns

These EEG and LFP results raised the following question: what is it in the brain activity of *rlss* mice that is different for such sleep-architecture and spectral variations to arise? Using MUA recordings and our recently developed ‘OFF-period’ detection algorithm (Harding et al., 2023), we discovered that neuronal firing in *rlss* mice ceases for seconds at a time during NREMS, in a locally synchronised manner (**Figure 2a and Movie 1**). Compared to WTs, in *rlss* animals, periods of such synchronised neuronal silence (‘OFF-periods’) lasted significantly longer and coincided with very low-amplitude, nearly isoelectric EEG and LFP traces (**Figure 2a,b and Movie 1**). Importantly, this also held true in REMS and wake when OFF-periods were generally longer in *rlss* mice (**Figure 2b**).

**Figure 2.**
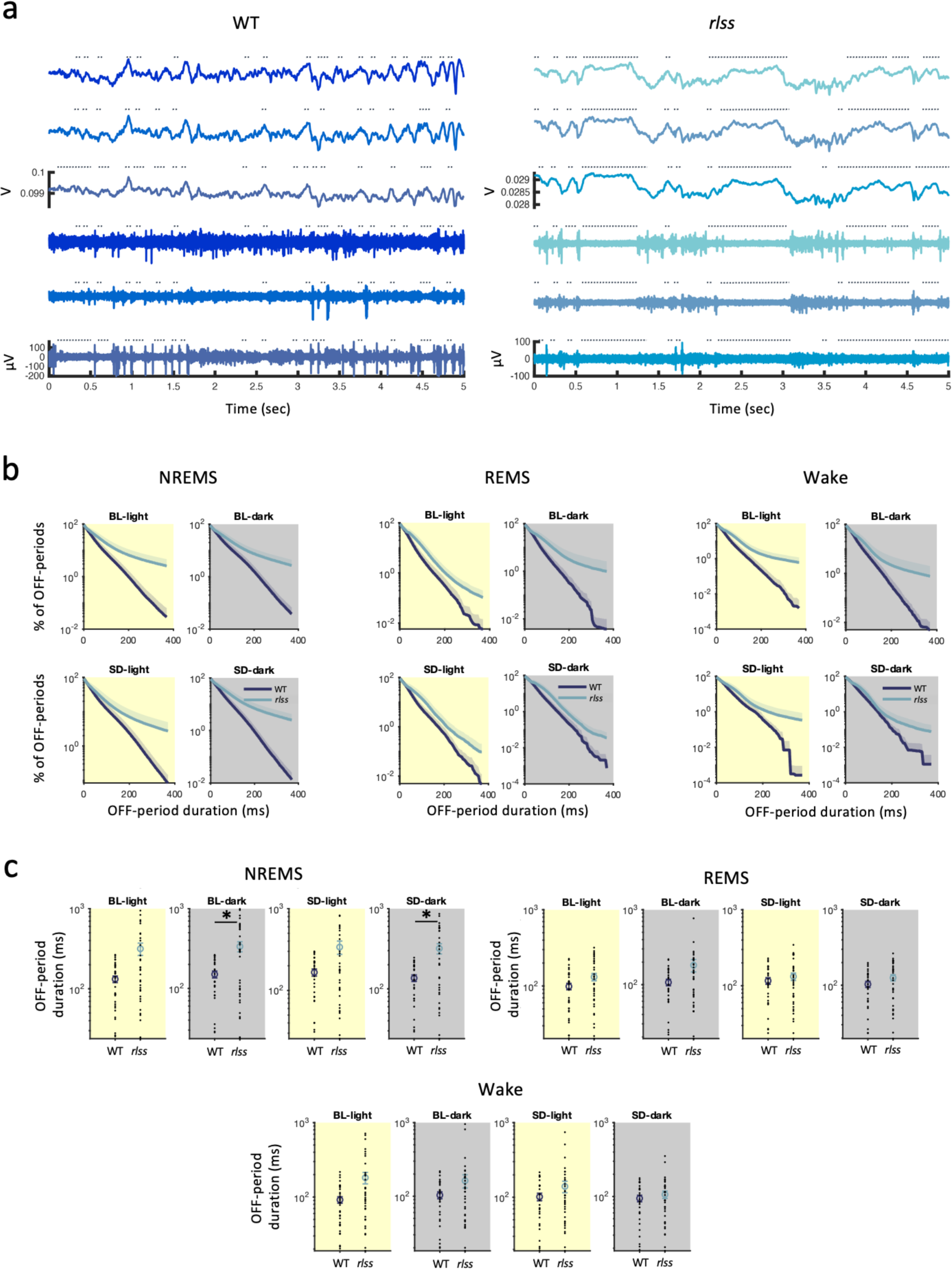
*rlss* mice show prolonged periods of neuronal silence across vigilance states that are reminiscent of burst suppression patterns. a) Examples of three LFP (upper traces) and corresponding MUA (lower traces) signals in a WT mouse (left) and *rlss* mouse (right) during 5 sec of NREMS. Dotted lines indicate location of OFF-periods as detected by the algorithm (see Methods) with no threshold applied. b) Survival curves of OFF-periods (relative to their duration) in NREMS (left), REMS (middle) and wake (right) during a baseline (BL) day (upper panels) and a SD day (lower panels), separately for the light and dark phases of a 24-h day in WT and *rlss* mice. Values shown as mean + SEM. c) Average OFF-period durations of the 5% longest OFF-periods during NREMS (upper panels), REMS (middle) and wake (lower panels), in a BL (left) or SD day (right) separately for the light and dark phases. Values shown as mean ± SEM, with black dots indicating individual channels. Wilcoxon rank sum test; * indicates p<0.05, **p<0.01, ***p<0.001. n(WT), n(*rlss*)=5. BL: baseline, SD: sleep deprivation.

As a next step, we aimed to specifically investigate the longest OFF-periods, and thus selected the 5% longest OFF-periods which we looked at separately (see Methods). We found that OFF-periods are significantly longer in *rlss* mice, being about twice as long as in WT during NREMS (**Figure 2c**); this further supports a state of ‘neuronal inertia’, whereby in *rlss* mice neurons are less likely to resume activity upon a period of silence of a certain duration.

As we previously hypothesised (Banks et al., 2020), it seems that changes in synaptic vesicular release lead to changes in cellular and network excitability which, by altering excitatory-inhibitory balance between cell populations controlling sleep and wake states, result in changes in sleep architecture. However, firing rates detected from MUA were not significantly altered (**Figure S1b**). This may also underlie the ‘default state’ features prominent in all vigilance states, leading to an increased uniformity in their spectral characteristics, particularly striking at state transitions (**Figure 1b,d,e**). We argue that transitions between active and silent states at the level of individual cells and networks may lag behind global state transitions, resulting in a longer time spent in mixed, hybrid states of vigilance.

Given this ‘neuronal’ inertia that may underlie the less clear-cut boundaries between vigilance state signatures, and the alterations in EEG and LFP spectra, notably in the slow-wave frequency range – which is traditionally used to investigate sleep homeostasis dynamics – we next asked whether sleep homeostasis was affected in *rlss* mice.

### Sleep homeostasis is generally conserved in *rlss* mice, with increased sleep quantity but not sleep quality, in response to sleep deprivation

To investigate the sleep homeostatic response in *rlss* mice we subjected the mice to a 6-h sleep deprivation protocol starting at light onset (time/Zeitgeber 0 - ZT0), after which the mice were left undisturbed for the subsequent 18h. Following this 6-h sleep deprivation period, *rlss* mice showed a markedly decreased sleep latency compared to their WT littermates (**Figure 3a**). This is the only case when we observed state switching probability being greater in *rlss* mice compared to WTs. The time spent in NREMS in the 6h following SD was increased by an 1h in *rlss* animals compared to WTs, while REMS time was halved (**Figures 3b-d**). However, across the whole recovery period (18h post sleep-deprivation), baseline comparative sleep structure was restored, with *rlss* mice spending less time in both NREMS and REMS than WT mice (**Figure 3d, right**). At the end of this 18-h recovery period, all mice had reached their baseline daily levels of NREMS and REMS (**Figures 3b,c**).

**Figure 3.**
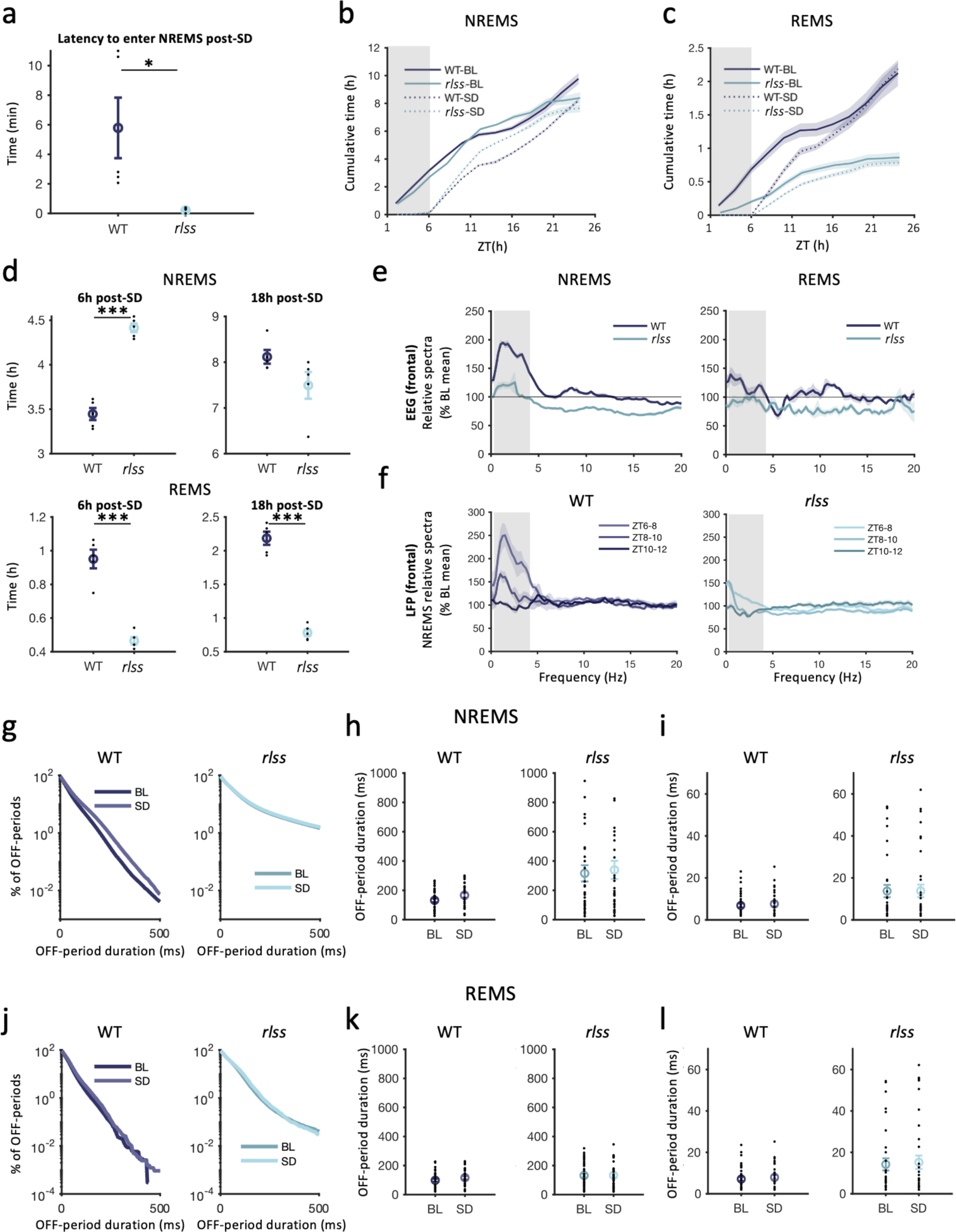
Response to sleep deprivation reveals a decreased sleep quality that is compensated for by an increase in sleep quantity in *rlss* mice. a) Latency to enter NREMS following a 6-h sleep deprivation (SD) protocol. b) Cumulative time spent (in 2-h bins) in NREMS in WT and *rlss* mice during a baseline (BL) or a SD day. Highlighted in grey is the SD period for the SD day only. c) As in b) for REMS. d) Amount of time spent in NREMS (top panels) or REMS (bottom panels) during the 6h (left panels) or 18h (right panels) following a 6-h SD protocol in WT and *rlss* mice. e) Frontal EEG spectra in NREMS (left) and REMS (right) relative to the baseline mean in WT and *rlss* during the 2h following a 6-h SD protocol. Delta frequency band (0.5-4 Hz) highlighted in grey. f) LFP spectra in NREMS relative to baseline mean in WT (left) and *rlss* (right) mice in the 6h following a 6-h SD protocol, separated in 2-h blocs. Delta frequency band (0.5-4 Hz) highlighted in grey. g) Survival curves of OFF-periods (relative to their duration) occurring during NREMS during the light phase of a BL and SD day in WT (left) and *rlss* (right) mice. h) Average OFF-period duration of the 5% longest OFF-periods in NREMS in the light phase of a BL and SD day in WT (left) and *rlss* (right) mice. Black dots indicate individual channels. i) As in h) but for the 5% shortest OFF-periods. j) As in g), but for OFF-periods occurring during REMS. k) As in h), but for OFF-periods occurring during REMS. l) As in i), but for OFF-periods occurring during REMS. Values shown as mean ± SEM, with black dots indicated individual values (mice in a & d, or channels in h, i, k & l). n(WT)=5 (3 for LFP-derived parameters), n(*rlss*)=5. Independent samples t-tests in a) and d); * indicates p<0.05, ***p<0.001. BL: baseline, SD: sleep deprivation, ZT: zeitgeber time.

SWA, the power in the delta (0.5-4Hz)-frequency range is a common marker of sleep need and sleep ‘efficiency’. Here, we found that the relative increase in SWA commonly triggered by sleep deprivation was markedly reduced in *rlss* mice (**Figures 3e and S2**). This was the case for both the EEG and LFP signals, and was particularly marked in the 2h directly following sleep deprivation (**Figures 3e,f and S2**). This indicates that although sleep homeostasis is partly conserved in *rlss* mice (the time taken to reach baseline levels of NREMS and REMS is comparable between genotypes – **Figure 3b,c**), SWA, reflecting sleep pressure, does not build up at the same rate in *rlss* mice. To summarise, *rlss* mice are able to compensate for sleep loss in a similar amount of time to WT mice (**Figures 3b,c**), but this is made possible by spending more time asleep (**Figure 3d**), as their increase in SWA during recovery NREMS (which allows for sleep pressure dissipation) is lower compared to their own baseline levels (**Figures 3e,f and S2**). In short, there seems to be a trade-off between sleep quantity (duration spent sleeping) and sleep quality (higher SWA levels) in those mice following a sleep deprivation challenge.

The fact that their baseline SWA levels is higher (possibly due to neuronal inertia) may also explain, firstly, why they sleep less overall during a baseline day, and secondly, why the relative increase in SWA following sleep deprivation is dampened. Indeed, there could be a ceiling effect with longer, more synchronised periods of neuronal silence not being possible as they are already long under baseline conditions. Therefore, a further increase in SWA (which reflects larger-amplitude, slow-frequency deflections in the signal and is thus a read-out of synchronized periods of neuronal silence and firing) may be limited. We observed that OFF-period durations following sleep deprivation did not increase in *rlss* mice, but nor did they in WT mice (**Figure 3h-i**). This can be expected as changes in the average duration of OFF-periods naturally occur between baseline and sleep deprivation conditions in WT animals (Vyazovskiy et al., 2009); therefore genotype-specific changes may be more easily observable in terms of incidence (**Figure 3g**) than duration. This may also be explained by the fact that here, we chose to analyse the longest OFF-periods and combine early and late sleep. Yet, early sleep usually contains longer OFF-periods than later sleep (McKillop et al., 2018; Vyazovskiy et al., 2009). No significant changes in OFF-period duration in REMS was observed in either WT or *rlss* mice in response to sleep deprivation (**Figure 3j-l**).

The fact that the increase in SWA is dampened in *rlss* mice compared to WT mice following sleep deprivation may either arise from *rlss* mice experiencing chronically elevated sleep pressure, thus leaving less capacity for a further increase in response to sleep deprivation. Or, it may be that as the neural activity of *rlss* mice persistently revolves around a default state, they do not accumulate sleep pressure as steeply as WT mice during waking (Thomas et al., 2020). To investigate this further, we used an adapted version of the ‘two-process model’ of sleep regulation that we have previously described (Guillaumin et al., 2018).

### Response to sleep deprivation in *rlss* mice reveals a decreased sleep quality that is compensated for by an increase in sleep quantity

To investigate the dynamics of SWA and sleep homeostasis in *rlss* mice, we applied the two-process model of sleep regulation (Achermann et al., 1993; Guillaumin et al., 2018) to recordings obtained in *rlss* and WT mice. This model predicts the levels of SWA across a 24-h day, given the hypnogram (vigilance states) of a mouse. We optimised the parameters of the model for each mouse against a baseline day (see Methods). The model’s predictions were satisfactory (squared error <10% at the end of the optimisation, see methods in Guillaumin et al. (2018)) but variable across mice (**Figure 4a**). In particular, the fit was poorer overall in WT mice (higher variations in the squared errors obtained at the end of the optimisation stage). This could stem from a more fragmented sleep-wake architecture in those animals, as opposed to *rlss* mice, which have more consolidated bouts of sleep and wakefulness due to their state inertia phenotype. As the model tends to perform better when faced with longer wake bouts (which present a ‘challenge’ to the model as they lead to higher subsequent SWA), we next performed another set of optimisations using 24 h of baseline followed by a sleep deprivation day (6 h sleep deprivation and 18 h recovery).

**Figure 4.**
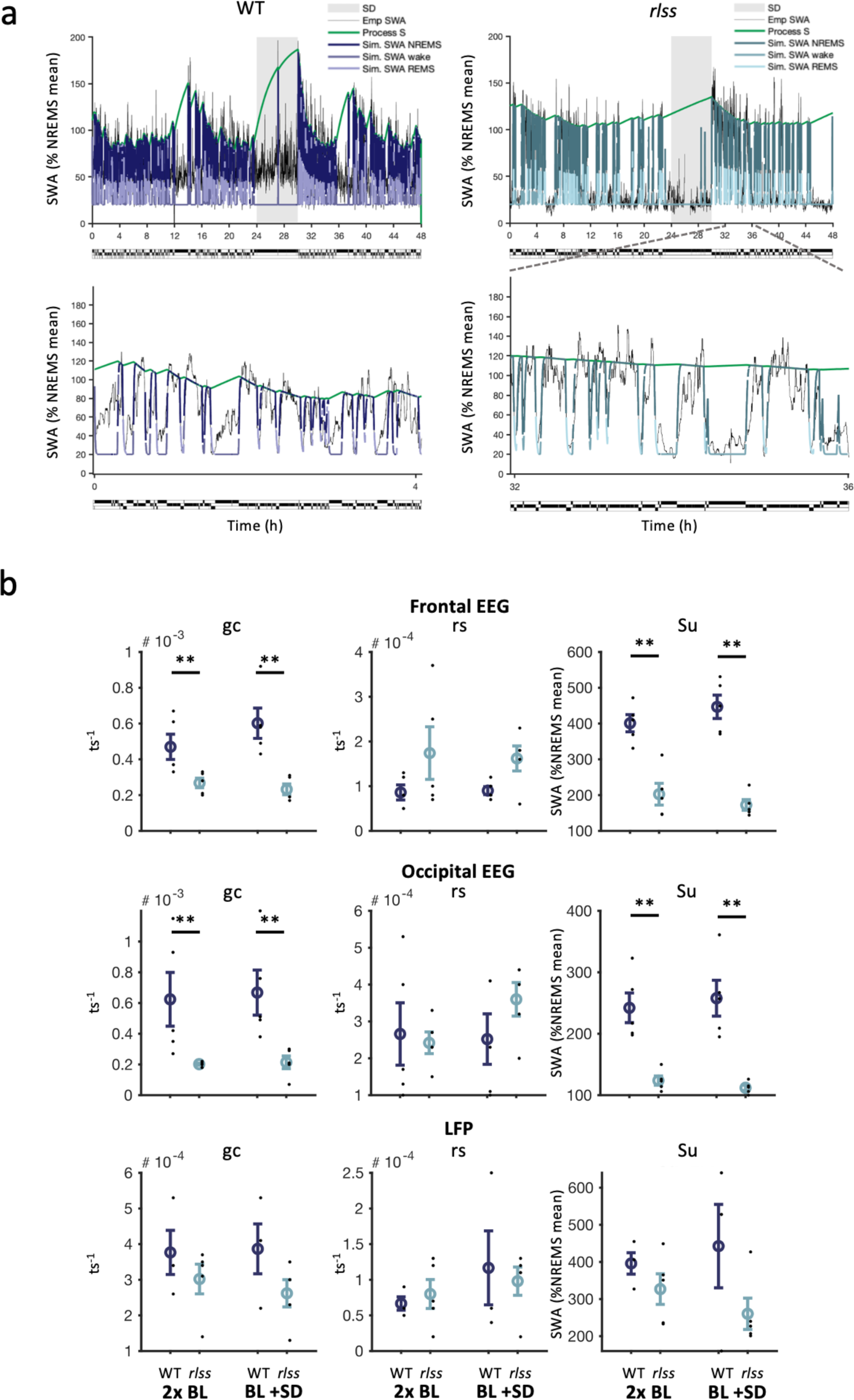
Sleep homeostasis dynamics are altered in *rlss* mice, with a lower decay rate of sleep pressure. a) Upper panels: Example of empirical and simulated slow-wave activity (SWA) across 48 h (24 h of baseline + 6-h sleep-deprivation + 18-h recovery) in an example WT (left) and *rlss* (right) mouse. Lower panels: zoomed-in inserts of a 4-hour section from the corresponding upper panels. Colours indicate the corresponding vigilance state, also shown underneath each panel as a hypnogram. b) Model parameters when modelling the SWA levels of the frontal EEG signal (upper panels), occipital EEG (middle panels) and LFP (lower panels), using 2 BL days (left panels of each parameter) or a BL+SD day (right panels of each parameter), in WT (n=5 for EEG, n=3 for LFP) and *rlss* mice (n=5). Values shown as mean ± SEM; black dots indicate individual values. Non-parametric Mann Whitney tests; ** p<0.01.

When comparing the results obtained with both approaches (BL only vs. BL+SD day), we found that the parameter values obtained with either approach were overall similar, except when optimisations were run against LFP data (**Figure 4b**). This suggests that the equations are better designed to fit EEG, from which they were initially derived (Guillaumin et al., 2018), than LFP signals. *rlss* mice had significantly reduced values of gc (gain constant, the rate of decrease of SWA) and S_U_ (upper asymptote of SWA levels) in both EEG derivations (**Figure 4b, upper and middle panels**), but not LFP (**Figure 4b, lower panels**). The results therefore suggest that the rate at which sleep pressure decreases in *rlss* mice (gc) is slower, but that the upper threshold to which sleep pressure can build-up (S_U_) is also lower. These findings should nevertheless be interpreted bearing in mind the model relies on power in the slow-wave range during NREMS (SWA), which is affected by the very low-amplitude oscillations observed in *rlss*, as described above.

Putting these results in perspective with our earlier findings, knowing that *rlss* mice have a reduced total sleep time, one could hypothesize a more efficient sleep, which would be reflected in a higher decay rate (higher gc), or hypothesize a slower build-up (lower rs) of sleep pressure during wake. Yet, we observed the opposite findings, with a significantly decreased decay rate in *rlss* mice compared to WTs, and no significant difference in the rise constant rs (**Figure 4b**). These results held true in both EEG derivations, but not in the LFP recordings. On the other hand, *rlss* mice were characterized by a significantly reduced upper asymptote S_U_. This could explain how *rlss* mice come back to baseline levels of sleep (**Figure 3b,c**) in the same amount of time as WT mice despite a slower dissipation of sleep pressure, given that they start with lower levels (as shown by a lower upper asymptote).

Taken together, computational modelling provided additional insight into the sleep homeostasis dynamics in *rlss* mice. Our data suggest that efficient sleep pressure dissipation requires intact neuronal dynamics, i.e. rate of shifting between quiescent and firing states, which is impaired in *rlss* mice. On the other hand, consolidated bouts of sleep, which *rlss* mice display, would not lead to better sleep pressure dissipation according to our model. In addition, these results support the idea that as *rlss* mice have intrusions of default-state/sleep-like patterns during wake (and REMS), they do not build sleep pressure as readily (Thomas et al., 2020). This is supported by our finding of a significantly reduced upper asymptote level, leading to a decreased need for sleep pressure dissipation.

### Conserved behavioural reaction to auditory stimuli, but altered response at the neuronal level, in ***rlss* mice.**

Finally, given the inertia shown by *rlss* mice to switch between states, both at the behavioural and neuronal levels, we asked whether their response to external stimuli – such as a sound – could be impaired too. We wanted to address whether the elevated power in the slow-wave range and the ceiling effect (smaller increase) in SWA following sleep deprivation could reflect a higher sleep depth in *rlss* mice, which would lead to a higher arousal threshold. Using an auditory stimulation paradigm (**Figure 5a**), we found that the number of full arousals (i.e. the mouse stays awake for at least 16 sec after the sound, see Methods) following the sounds was similar across genotypes, but there were significantly fewer brief arousals (<16 sec) in *rlss* mice (**Figure 5b**). This may reflect once again the state inertia previously described in those mice (Banks et al., 2020), whereby once *rlss* mice have entered a state they tend to remain in that state for longer, hence the lower likelihood of a brief arousal (if the mouse does wake up, then it stays in that state longer as it cannot switch back to NREMS as easily). This hypothesis is further supported by the occurrence of brief awakenings (short arousals ≤16 sec) during NREMS, which is three times less likely in *rlss* mice during undisturbed (baseline) recordings (**Figure 5c**). Of note, the number of muscle twitches in response to the sound was similar across mice (F(1,7)=0.002, p=0.967, **Figure 5b, right**), indicating no obvious impairment in acute motor or EEG response to environmental triggers (**Figure 5e**).

**Figure 5.**
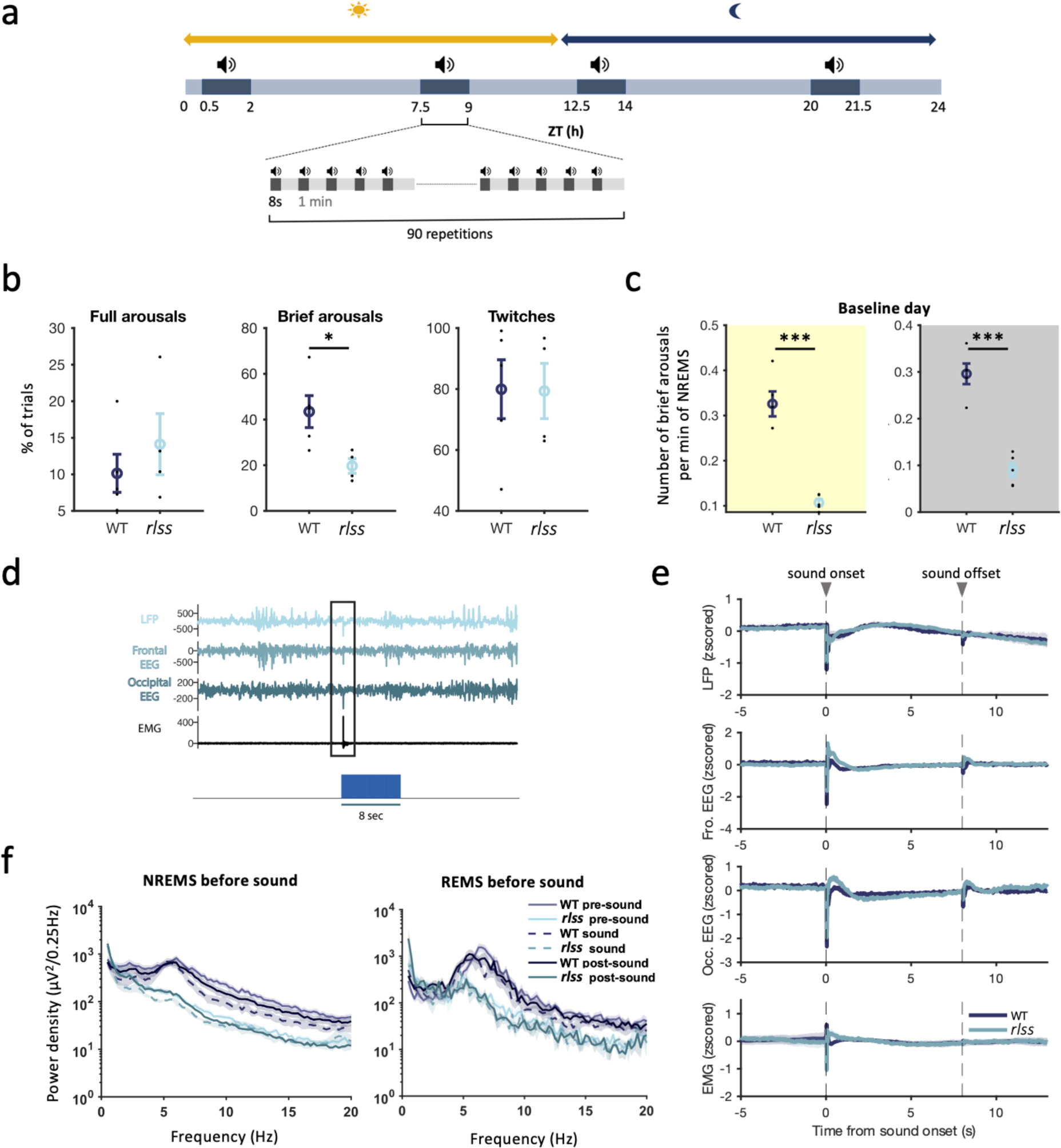
*rlss* mice show a conserved arousal threshold but an impaired neuronal response to an aurditory stimulus during sleep. a) Schematic of experimental design. b) Number of full arousals, brief arousals and muscle twitches in response to a sound. c) Number of brief arousals per min of NREMS during the light and the dark phase of a baseline day. d) Example of a muscle twitch and associated artifacts in the LFP and EEG channels in response to a sound. e) LFP (1 representative channel per mouse), frontal EEG, occipital EEG and EMG around sounds. Signals are extracted from all sound trials occurring during the light phase, and z-scored before being averaged across trials and across mice. f) Power density spectra calculated from the occipital EEG in the 8 sec before, 8 sec during and 8 sec after a sound, if the sound onset occurred during NREMS (left) or REMS (right). n(WT)=5 (3 for LFP), n(*rlss*)=4. Values shown as mean ± SEM; black dots indicate individual values. Independent samples t-tests; *p<0.05, ***p<0.001. Occ.: occipital, ZT: zeitgeber time.

When looking at the EEG spectra during the sound (8 sec) and during segments of the same length either before or after the sound, we found that – as expected given baseline signatures – there was a significant difference between genotypes. More strikingly, there was a significant Time Point x Genotype interaction if the mouse was in NREMS when the sound occurred (F(2,14)=5.431, p=0.018), but not if the mouse was in REMS when the sound started (F(2,14)=1.395, p=0.280) (**Figure 5f**). This indicates that if a sound occurs while a mouse is in NREMS, the evolution of the response in terms of electrophysiological signature from pre-until post-sound epochs is significantly different between WT and *rlss* mice. More specifically, the change in the EEG spectra in response to the sound was more pronounced in WT mice than in *rlss* mice when the mice were originally in NREMS before being woken up.

Thus, *rlss* mice show a conserved arousal threshold, as indicated by the comparable number of full arousals and twitches in both genotypes. Yet, similarly to their response to sleep deprivation, it appears that *rlss* mice show a ‘rigidity’ in their neuronal dynamics, so that the scope for variation (as measured by spectral power) across states and wake/sleep experiences is markedly reduced.

## DISCUSSION

*rlss* mice are characterized by a mutation in the VAMP2 SNARE protein, that induces a change at the molecular scale by impairing neurotransmitter release (Banks et al., 2020). Here, we found that *rlss* mice display a state inertia, switching less between states, both at behavioural (vigilance state) and at cellular (neuronal) scales. Our results therefore support the idea that sleep regulation is tightly linked to synaptic function. In particular, we show that *rlss* mice display long periods of neuronal silence, strikingly reminiscent of deep anaesthesia, and which underlie increased SWA during baseline conditions, but lead to an inability to fully adapt neural dynamics in response to behavioural experiences such as sleep deprivation or auditory stimuli. Mathematical modelling revealed that the rate of sleep pressure dissipation is slower in those mice. Finally, we found this neural rigidity is compensated for by global behavioural states, such as increased sleep time in response to sleep deprivation.

### An impairment in the ‘sleep switch’?

Our previous findings (Banks et al., 2020), combined with the auditory stimulation protocol results described here support the idea of ‘vigilance state inertia’ in *rlss* mice. The decrease in neurotransmitter release efficiency leads to a difficulty in switching between two consecutive vigilance states in those mice. Previous studies have suggested a mechanism for a sleep/wake switch in both flies and rodents (Pimentel et al., 2016; Saper et al., 2010). ‘Flip-flop’ switches have been proposed, either to control the transitions between REMS and NREMS (Lu et al., 2006), or to allow rapid transitions between NREMS and wake and prevent animals from lingering between two vigilance states – half awake and half asleep (Saper et al., 2001; Saper et al., 2010). Saper et al. (2001) propose that if neuronal firing is weakened on either side of this bistable switch, then the stability of the switch is compromised; on the other hand, they also propose that the time it takes to transition from one state to the next may depend on the ability for a population of neurons that promote a given state to overcome the ‘resistance’ of the population of neurons promoting the current state (Saper et al., 2010). In *rlss* mice, we observe an increased stability of each state, both at the behavioural and cellular levels. Concomitantly, we see more similar spectral signatures between vigilance states. This indicates that not only is the sleep-switch not transitioning between states as readily in those mice, but also that it is ‘leaky’, staying halfway between states, resulting in mixed states with characteristics of both sleep and wake.

Another mouse line (*Sleepy*), developed using a forward genetics approach, exhibited increased NREMS time; the gain-of-function mutation present in these mice affected the *Sik3* gene, and led to a shorter SIK3 protein kinase, which is likely involved in the regulation of sleep amount via its phosphorylation status (Funato et al., 2016). Interestingly, despite this hypersomnia, in contrast to the baseline behavioural phenotype of *rlss* mice, the NREMS delta power was increased in *Sleepy* mice, just like in *rlss* mice. *Sleepy* mice showed an exaggerated response to sleep deprivation, suggesting an increased inherent sleep need despite an increased delta power during baseline wake. The ability of *Sleepy* mice to stay awake when exposed to arousal-promoting stimuli was conserved, showing that, as with *rlss* mice, their ability to be awake should the need arise is not impaired (Funato et al., 2016). But their accumulation of sleep pressure was enhanced leading to longer sleep time when left in undisturbed conditions.

Another mouse model obtained with a forward genetics approach displayed decreased REMS time, with REMS episodes lasting for shorter durations (Funato et al., 2016). The authors thus named those mice *Dreamless* and proposed that the mutation present in this new line (a gain-of-function mutation in the sodium leak channel *Nalcn* gene) led to an increased excitability of REMS-inhibiting neurons. In other words, one could hypothesize that an increase in excitability of REMS-inhibiting neurons would destabilise the sleep-wake switch, leading to more frequent REMS-to-wake/NREMS transitions, and to shorter REMS duration due to the instability of the REMS state.

These two other examples, which display some overlapping behavioural phenotypes to the *rlss* mice, do not support the idea that specific genes directly regulate sleep-wake architecture. Rather, those examples back the hypothesis that synaptic function and neuronal excitability – both of which alter the fine balance in neural networks cross-talks – are crucial for sleep-wake regulation.

An important point to be discussed, however, is whether the decreased state-transitioning probability observed in *rlss* mice signifies that it is uniformly difficult for them to switch to another state, or rather, that one state is at a stronger disadvantage. In particular, because of the NREMS-like features of wake and REMS in *rlss* mice (e.g. lower power in higher frequencies), it may be less relevant for them to enter full NREMS. We would thus observe an absolute decrease in the probability to enter NREMS relative to other states. And indeed, this is supported by what we observed when looking at state transitioning probabilities from Banks et al. (2020): there is an absolute 44% change in probability to transition between wake and NREMS (wake->NREMS: 34% decrease, NREMS->wake: 10% increase) and a 16% change in probability to transition from REMS to NREMS (REMS->NREMS: 87% decrease, NREMS->REMS: 71% decrease). The REMS->wake transitions are relatively less affected (absolute 8% decrease). This would support the idea that rather than a ‘state inertia’, those mice revolve around a ‘default state’ mode (Bandarabadi et al., 2020; Buzsáki, 2006; Hinard et al., 2012; Lemieux et al., 2014; Sanchez-Vives et al., 2017; Sanchez-Vives and Mattia, 2014; Timofeev et al., 2020) regardless of their vigilance state, thus making it less relevant to spend time in NREMS.

### Higher sleep pressure or a higher propensity to return to a ‘default state’?

The shorter sleep time in baseline conditions observed in *rlss* mice raises the question of whether these mice have higher sleep efficiency (they can dissipate sleep pressure faster), a decreased sleep pressure build up during waking or that they live under elevated sleep pressure. The two-process model of sleep regulation, which predicts the evolution of Process S, is a useful tool to address this question. A study in humans comparing short sleepers and long sleepers found that both groups had the same constants of increase and decrease of process S, which implied that short sleepers may live under a higher sleep pressure (Aeschbach et al., 1996). Here, we find the *rlss* mice have significantly lower rates of decrease but also a significantly lower upper asymptote of S. These results may therefore indicate than instead of having an inherently elevated sleep pressure (which their lower S upper asymptote argues against), *rlss* mice have a higher propensity to enter or approach the ‘default state’, due to increased inhibitory tone and/or decreased excitatory tone (induced by the slower neurotransmitter release).

There is evidence that the cortical oscillations are not merely an epiphenomenon, or simply a substrate for sleep-related processes to occur (e.g. memory consolidation), but have a role in vigilance state control and sleep regulation (Krone et al., 2021). Thus, if the cortex keeps track of the accumulation of sleep need, then the persistent sleep-like patterns we observe in our cortical recordings could indicate a partial sleep debt discharge across both sleep and waking states.

### Impaired vesicular release leads to longer neuronal OFF-periods

Our MUA recordings revealed striking neuronal firing patterns in *rlss* mice during NREMS. Slow-waves were associated with prolonged periods of neuronal silence, where neighbouring neurons stop firing in a coordinated manner for prolonged periods of time, with the longest OFF-periods being about twice as long in *rlss* mice as in WT mice (**Figure 2**). Those periods of neuronal silence resulted in very-low amplitude LFP signals. Such patterns of neuronal activity are seldom seen during natural sleep and are a more common feature of general anaesthesia (Antkowiak, 2002; Lukatch and MacIver, 1996).

We have previously described an impairment in vesicular release in *rlss* mice due to the mutation in *Vamp2*, which plays a key role in vesicular and cellular membrane fusion (Banks et al., 2020). *rlss* mice showed a marked decrease in vesicular-release probability following single action potentials. An impairment in vesicular release could lead to a state akin to anaesthesia. Indeed, it has been shown that general anaesthetics, such as isoflurane, propofol or etomidate, induce a decrease in synaptic neurotransmitter release machinery by interacting with multiple SNARE proteins (Herring et al., 2009; Xie et al., 2013). This could account for the long periods of neuronal silence observed in *rlss* mice while they are (naturally) asleep: given their mutation leading to a structural change in VAMP2, which is a SNARE protein, *rlss* mice may present features of an ‘inherently anesthetised’ state.

Interestingly, it was previously shown that injection of the GABA_A_ agonist muscimol led to a catalepsy-like state, followed by a hyperactive state in mice, with the EEG of both of those states displaying increased power in the slower-frequency range at the expense of the higher-frequency range (Vyazovskiy et al., 2007). This is strikingly similar to our findings, with increased synchrony in all states, and reduced power in high frequencies, alongside a hyperactive phenotype. Furthermore, the dissipation of sleep pressure did not occur under muscimol despite its SWA-enhancing effect, which, the authors conclude, suggests a role for normal physiological sleep in dissipating sleep pressure (Vyazovskiy et al., 2007). Again, this parallels our findings in *rlss* mice which have a markedly reduced constant of decrease of SWA (**Figure 4b**). The authors propose that the effects of muscimol point towards a crucial role for inhibitory connections in slow wave synchronization. Thus, our results suggest that the decreased vesicular release is in many ways akin to increased inhibitory tone and leads to alterations in inhibitory-excitatory balance. The physiological sleep of *rlss* mice is in several ways more similar to non-physiological states such as cataplexy or anaesthesia (e.g. with an inefficiency to properly dissipate sleep pressure) than normal sleep.

### Switching between anaesthesia and wake

Studies in Drosophila have shown that some ‘narrow abdomen’ mutants with an altered stability in the switch between wakefulness and sleep displayed increased sensitivity to anaesthesia (Joiner et al., 2013; Krishnan and Nash, 1990). Those mutants present with an altered response to halothane (Krishnan and Nash, 1990) and isoflurane (Joiner et al., 2013): the flies lost consciousness in less time than the WT flies, but were more resistant to the anaesthetic’s effects on motor control (Krishnan and Nash, 1990). It was shown that those narrow abdomen mutant flies have altered transitions between wake and anaesthesia, and the authors support the view that the narrow-abdomen gene is part of 4 genes necessary for ‘neural inertia’, allowing mutual inhibition between arousal-promoting and arousal-suppressing loci, forming a bistable switch between waking and anesthetized states, similar to the sleep switch (Joiner et al., 2013). And indeed, the mutation in those flies affects the orthologue gene to the one affected in the Dreamless mice mentioned above, that present with decreased REMS-state stability. A later study went on to show that the mutation affecting the narrow-abdomen flies impacts neuronal excitability and firing rates under the control of the circadian clock, and participated in a ‘bicycle model’ of membrane excitability (Flourakis et al., 2015). This seems to further support the idea discussed above that switching between states requires tightly-regulated excitatory and inhibitory balance between populations of neurons. Also, as mutations with similar effects on neural inertia but opposite effects on anaesthesia induction were observed in flies, it was proposed that the genes (but also possibly brain regions) responsible for anaesthesia induction and state stability are distinct (Joiner et al., 2013). The authors also proposed that the molecular and anatomical pathways involved in sleep homeostasis also contribute to neural inertia (Joiner et al., 2013). This would be in line with our findings, whereby state switching is impaired, and yet, when sleep pressure is high, mice are able to quickly shift to NREMS. Additional evidence suggests that the reversible loss of consciousness caused by anaesthetics occurs via the arousal and sleep pathways, involving both the thalamus and hypothalamus (Franks, 2008), and that the mechanisms underlying sleep homeostasis and anaesthesia overlap (Baker et al., 2014; Gelegen et al., 2018; Pal et al., 2011; Steinberg et al., 2015; Tung et al., 2002).

Yet, these observations start from the premise that sleep and wake are mutually exclusive states between which animals switch in a clear-cut manner, which is now acknowledged to be more complex (Nir and de Lecea, 2023). Indeed, this may not always be the case, as clearly shown by several studies (Bréant et al., 2022; Jang et al., 2022; Vyazovskiy et al., 2007), and states may indeed be viewed on a continuum, with some feature overlap. This is indeed what we see in *rlss* mice with an intrusion of NREMS-like features in REMS and wake states. It is remarkable that the *rlss* mice display only mildly impaired cognitive and behavioural phenotypes typical of wakefulness (Banks et al., 2020), despite the profound alterations in brain activity observed in those mice. Just as the occurrence of slow-waves or decreased neuronal activity which have been previously described in waking and not only in sleep (Fisher et al., 2016; Vanderwolf and Robinson, 1981; Vyazovskiy and Tobler, 2012), our findings further support that vigilance states are not necessarily separate phenomena with unambiguous boundaries, but can share characteristics. Such overlapping features may thus fluctuate depending on specific waking behaviour (Fisher et al., 2016), chemical manipulations (Bréant et al., 2022) or genetic alterations (Funato et al., 2016).

### Conclusion

This combination of *in vivo* and computational work provides new insights into the mechanisms that underlie sleep architecture and the alternation between vigilance states, as well as the generation of slow waves. Our results support the idea that synaptic function, and thus neuronal excitability, are direct contributors to sleep-wake architecture. The preserved ability of *rlss* mice to respond to salient external stimuli suggests that neuronal dynamics and sleep-wake architecture can be markedly altered before this leads to a phenotype that would represent an evolutionary disadvantage, with the animal less likely to survive natural selection. Ultimately, our findings regarding the mixed spectral signatures across states in *rlss* mice further support the idea that wake and sleep are not homogeneous states, but rather stand on a continuum of many possible states between full wakefulness and anaesthesia/loss of consciousness.

## METHODS

### Animals and housing conditions

All work was carried out in accordance with the UK Animal [Scientific Procedures] Act 1986 and the University of Oxford’s Policy on the Use of Animals in Scientific Research, and in compliance with the Animal Research: Reporting In Vivo Experiments (ARRIVE) guidelines. Mice from the *rlss* colony (MGI:5792085, C3H.Pde6b+ background) were initially obtained from MRC Harwell (U.K.) and were then bred in-house. N=5 *rlss* homozygotes and n=5 WT littermates (all males) were implanted. The experimenter was not blind to the animals’ genotype. Over the two months preceding surgeries, to promote weight gain, *rlss* homozygote mice were fed a high-nutrient diet (RM3 diet, ground, Special Diets Services, U.K.) in the form of mash in addition to the regular pellets. Mice were implanted once they had reached a weight nearing or above 30g, which meant that mice were overall 9 weeks older at surgery. The average age and weight of the animals at the time of surgery was 23.8 ± 1.2 weeks & 31.9 ± 0.8 g for the homozygotes and 14.7 ± 1.5 weeks & 31.7 ± 1.5 g for the WT mice. Age differences were permitted so that animals were matched by weight. Previous data indicates that this age difference does not contribute to the differences in sleep observed (Banks et al., 2016; McKillop et al., 2018).

### EEG and laminar probe implantation surgeries

On the day preceding surgery, mice were injected with Dexamethasone (0.2 mg/kg, subcutaneous route). Surgeries were performed using aseptic techniques under isoflurane anaesthesia (1-2%). Mice were provided with meloxicam (5 mg/kg s.c.), buprenorphine (0.1 mg/kg) and Dexamethasone (0.2 mg/kg) prior to surgery start. Screws (self-tapping bone screws, stainless steel, length: 4 mm, shaft diameter: 0.85 mm, Fine Science Tools Inc.) were positioned (Motor cortical area (frontal): AP +2 mm, ML +2 mm; Visual cortical area (occipital): AP −3.5 mm, ML + 2.5 mm; Cerebellum: 2 mm behind lambda; Anchor screw: AP −3.5 mm, ML −2.5 mm). Frontal EEG signals are well suited to measure SWA and detect spindles, while occipital EEG signals are better suited to investigate theta activity, active wakefulness and REMS. A small craniotomy was performed around the laminar probe (type A1×16-3mm-100-703-Z16, NeuroNexus Technologies, Ann Arbor, USA) insertion point (≈0.8mm x 0.8 mm, primary motor cortex, AP +1.1, ML −1.75, DV 1.8, tilt of 14°). The tip of the electrode was dyed prior to the surgery with 1,1’-Dioctadecyl-3,3,3’,3’-Tetramethylindocarbocyanine Perchlorate dye (‘DiI’, ThermoFisher Scientific). The probe was grounded to the anchor screw and referenced to the cerebellar screw and carefully lowered into the cortex (1.8 mm from the point at which the tip touches the surface of the brain). Silicon gel (Kwik-Sil, World Precision Instruments, USA) was applied on the craniotomy and the implant cemented. Two electromyography (EMG) electrodes were inserted in the neck muscle and fixed to the back of the head cap with dental cement. Saline was injected subcutaneously (0.1 mL/20 g) to compensate for fluid loss during the duration of the surgery, before placing the mouse in a recovery chamber at 28°C. Animals were administered with Dexamethasone (0.2 mg/kg, s.c.) and meloxicam (5 mg/kg, s.c.) for 1 and 3 days following surgery, respectively.

### Recordings and Initial Data processing

Animals were left to recover for at least 2 weeks before being moved to custom-made clear plexiglas cages (20.3 x 32 x 35 cm^3^) placed in ventilated, sound-attenuated Faraday chambers (Campden Instruments, UK). Mice were kept in a 12h:12h light-dark cycle (lights on at 9am), with water and food available ad libitum. Temperature within the room was kept at 22±2 °C, with a humidity level of 55±10 %, and light levels of 120-180 lux.

Mice were left to habituate to their new environment for 3 days before being tethered. Recordings were started at least 3 days after tethering, to let the mice habituate to the cables. Signals were recorded using the Multichannel Neurophysiology Recording System (TDT, USA) and the Synapse software (TDT, USA). As described earlier (Banks et al., 2020), EEG and EMG signals were filtered (0.1-100 Hz), amplified (PZ5 NeuroDigitizer preamplifier, Tucker-Davis Technologies) and stored at a sampling rate of 256.9 Hz. Signals were extracted and converted using custom-written Matlab (The MathWorks Inc., USA) scripts as previously described (Guillaumin et al., 2018; McKillop et al., 2018). Vigilance states – wake, NREMS, REMS, or brief awakenings (short arousals ≤16 sec during sleep, as described previously (Vyazovskiy et al., 2009)) – were scored manually using the SleepSign software (Kissei Comtec Co, Nagano, Japan). EEG power density spectra were computed by a Fast Fourier Transform (Hanning window; 0.25-Hz resolution). More detailed recording and initial analysis methods were as published (Banks et al., 2016; Banks et al., 2020; McKillop et al., 2018). As the focus of the present study was on the slower frequencies, spectra only show power for frequencies up to 20 Hz.

### Sleep deprivation

Sleep deprivation (SD) was performed for 6 consecutive hours starting from light onset (ZT0). Animals were kept awake by providing them with new objects to explore. If the mice were getting drowsy, nesting material was taken away from the cage temporarily. SD was successful as only 7.8 ± 1.7 minutes (mean ± SEM) of sleep were detected during the 6 h of SD. Following sleep deprivation, mice were left undisturbed for at least 18 h.

### Auditory stimulation

Sounds of 8kHz, 75dB were played for 8 sec every minute during 1h30 (90 repetitions of 8-sec sound, 52-sec silence). This 1h30 session was performed 4 times across 24 h: ZT 0.5 – 2; ZT 7.5 – 9; ZT 12.5 – 14; ZT 20 – 21.5. Speakers were placed 70 cm above each animal, within the Faraday sound-attenuated chambers.

### Detection of OFF-periods

Detection of low-amplitude neural activity segments (‘OFF-periods’) was performed using the GUI made available by C.Harding, described in Harding et al. (2023). Default parameters of the GUI were used for OFF-period detection (Calinski-Harabaz clustering evaluation method; Clustering variable 1: amplitude, 15 smoothing points; clustering variable 2: amplitude, 5 smoothing points; clustering sample size: 1%). Out of the 16 channels, 5-10 MUA channels (and corresponding LFP channel for visualization) were selected for each mouse, selecting only those that were stable across a 48h (baseline day + sleep deprivation day). The first 12 h of the baseline day were used as a new dataset against which the detection parameters were calculated for each mouse. The parameters (‘pre-sets’) were then re-used to detect OFF-periods for the next 36 h of recording.

In some instances (specified in the legends and text), only the 5% longest OFF-periods were included. The 5% threshold was chosen to include enough OFF-periods, while still focusing on the longest ones, wherever that was the focus of our investigations.

### Application of the ‘elaborated two-process model’

The estimation and optimisation of the model parameters (gain constant gc, describing the decay rate of SWA; rise constant rs and upper asymptote S_U_) were performed exactly as described in Guillaumin et al. (2018), using SWA calculated across a 24-h baseline day and a sleep deprivation day (6-h sleep deprivation + 18-h recovery sleep). For the baseline-only optimisations, the baseline recordings were artificially duplicated for the estimation/optimisation of the parameters, as described in the above-mentioned publication. For the BL+SD day, the 24-h baseline day followed by the sleep deprivation day were used.

### Perfusions and micro-lesions

Mice were injected with Pentobarbital (200mg in 1 mL, i.p. injection of 0.01mL/g body weight). Microlesions of 3 selected channels on the laminar probe were then performed using a NanoZ device (White Matter) before the mice were transcardially perfused with a PBS solution, followed by paraformaldehyde (PFA, 4% in PBS). Brains were removed and placed in PFA (4% in PBS) for 48 h, after which they were transferred to a 30% sucrose solution (in PBS) for cryoprotection. Once sunk to the bottom of the vial, brains were briefly rinsed in PBS and embedded in O.C.T. compound and frozen at −80°C until cutting.

### Histology

To confirm electrode placement, slices were prepared using a cryostat (Cryotome FSE, ThermoScientific; −22°C, 50μm-thick sections) and mounted on glass slides. Slides were subsequently mounted under cover glasses with ProLong Glass Antifade mountant with NucBlue stain (ThermoFisher Scientific). Sections were digitized using an LSM 710 laser scanning confocal microscope and ZEN image acquisition software (Zeiss) (**Figure S3**).

### Statistical analysis

All statistical analyses were performed using SPSS (IBM Corp., Release 23.0.0.2) and Matlab (The MathWorks Inc., USA). The nature of the tests used and corresponding results are reported within figure legends or within the main text of the Results section. Wilcoxon rank sum test was used where normality was not observed (Kolmogorov Smirnov test).

## Acknowledgements

We thank Prof. Peter Achermann for advice on Process S simulations and Prof. Irene Tracey for stimulating discussions and valuable suggestions regarding data interpretation. We also thank Dr Laura E. McKillop for valuable advice on surgical procedures. M.C.C.G. was supported by a BBSRC DTP grant (BB/J014427/1), a Swiss National Science Foundation grant (no. 310030_189110) and a Clarendon Fund Scholarship from the University of Oxford. C.D.H. was supported by funding from the Engineering and Physical Sciences Research Council (EPSRC, EP/S515541/1). L.B.K. was supported by a Wellcome Trust PhD studentship (203971/Z/16/Z) and by a Mann Senior Scholarship in medical sciences at Hertford College, Oxford. T.Y. was supported by the Naito Foundation (Grant for Studying Overseas), a Uehara Memorial Foundation (Postdoctoral Fellowships for Research Abroad) and a Wellcome Trust Senior Investigator Award (106174/Z/14/Z). M.C.K. was supported by a Berrow Foundation Lord Florey Scholarship. C.B.-D. was supported by a Wellcome Trust PhD studentship (109059/Z/15/Z) and a Clarendon Fund Scholarship from the University of Oxford. P.M.N. was supported by the MRC (grant codes MC_U142684173 and MC_UP_1503/2). S.N.P. was funded by the BBSRC (BB/ I021086/1) and a Wellcome Trust Strategic Award (098461/Z/12/Z). V.V.V. was funded by a John Fell OUP Research Fund Grant (131/032), a Wellcome Trust Strategic Award (098461/Z/12/Z) and by Medical Research Council (UK) Grant MR/S01134X/1.

## Author contributions

MCCG: implantation surgeries, experiments and data collection, data analysis, writing of the manuscript and figures, funding support.

CH developed the OFF-period detection algorithm.

LBK contributed to the development of the OFF-period detection algorithm and advised on implantation, microlesions and histology.

TY advised on surgeries and auditory stimulation protocol. MCK advised on technical issues.

CBD advised on methodology and data analysis for the auditory stimulation experiment.

GTB and PN assisted in the development of the mouse model and initial phenotyping of the *rlss* mice. SNP advised on study design, methodology, supervision, funding support.

VVV: conceptualisation/study design, methodology, supervision, direction of data analysis and manuscript preparation, funding support.

## SUPPLEMENTARY FIGURE PANELS

**Figure S1.**
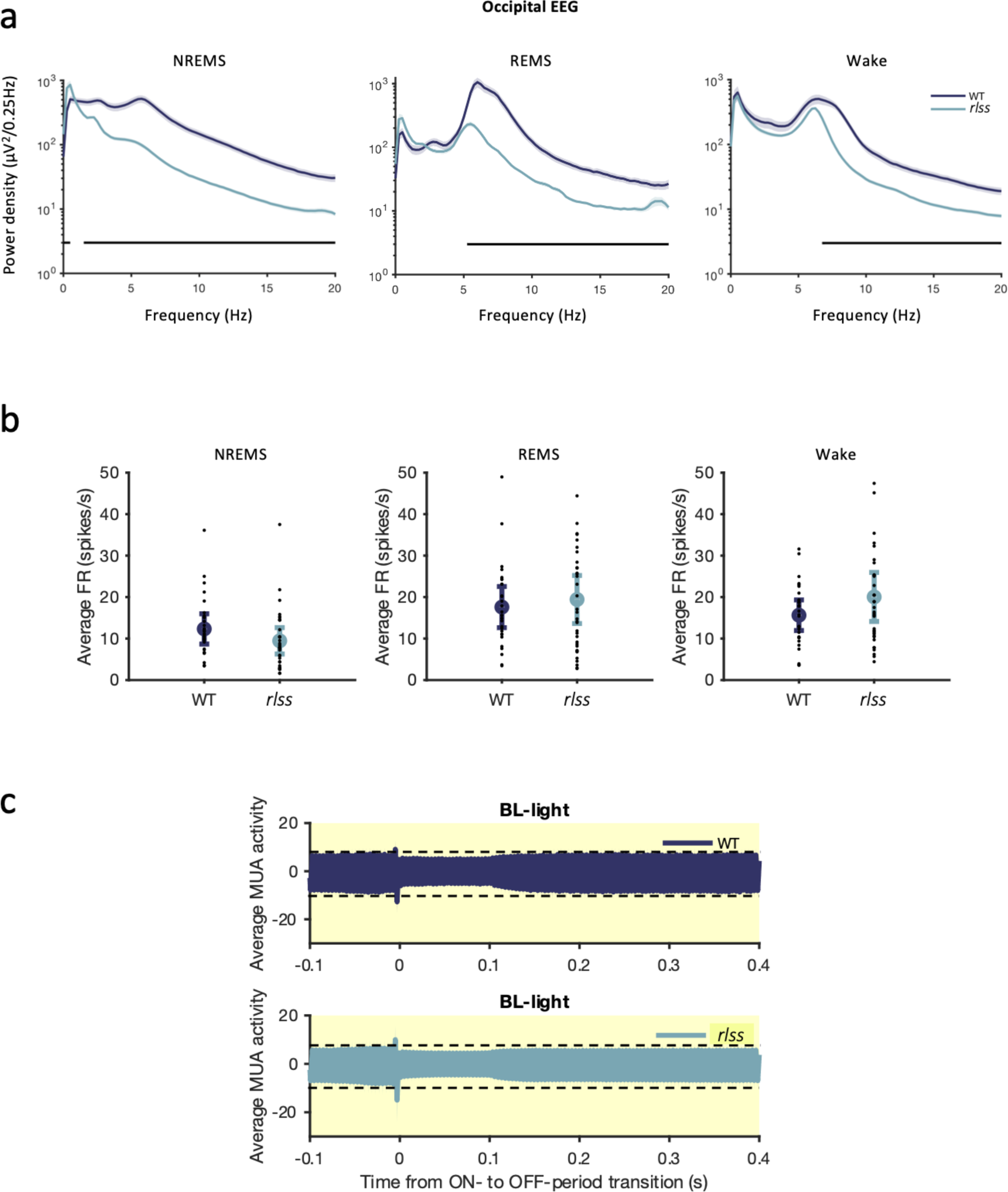
a) Occipital EEG power spectra during NREMS, REMS and Wake in WT and *rlss* mice. Black bars at the bottom indicate the frequency bins for which the power is significantly different (p<0.05) between WT and *rlss* mice. b) Average firing rate (FR, calculated from the multi-unit activity (MUA) as number of spikes per second) in each vigilance state, calculated from the light phase of a baseline day. n(WT)=4, n(*rlss*)=5. c) Average multi-unit activity across channels at ON-to OFF-period transitions in WT mice (top) and *rlss* mice (bottom). Only transitions during a 12-h light phase of a baseline day is included and only transition to OFF-periods that last at least 100 ms were considered here.

**Figure S2.**
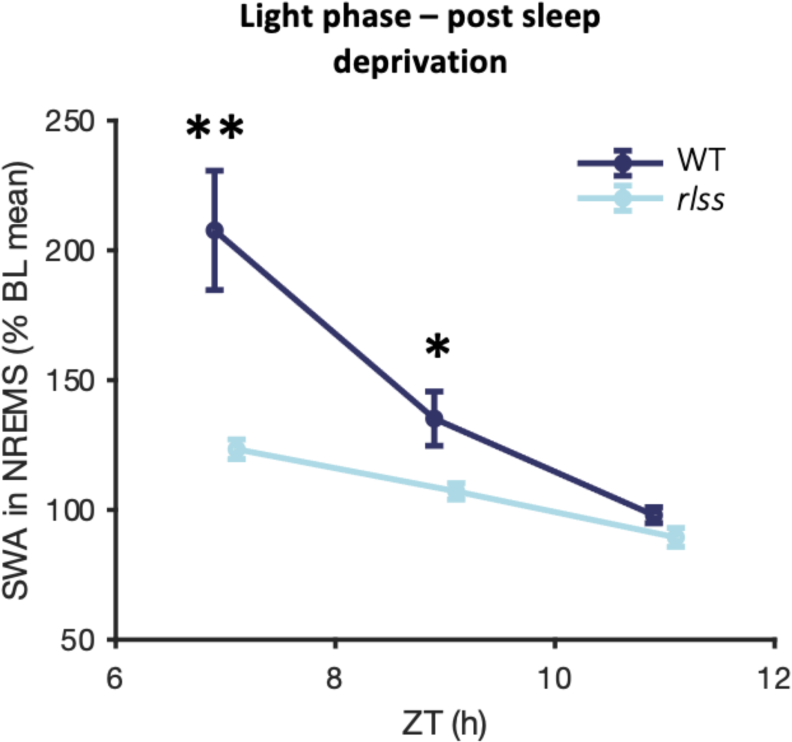
SWA levels during NREMS, from LFP recordings, in the 6 h following 6 h of sleep deprivation. Values normalised against a 24-h baseline mean, plotted in 2-h bins. Mean ± SEM; n(WT)=3, n(Hom)=5. Two-way ANOVA revealed a main effect of Time Point (F(2,12)=80.604, p<0.001), Genotype (F(1,6)=17.313, p=0.006), and a significant interaction Time Point* Genotype (F(2,12)=23.745, p<0.001). Post-hoc testing with Bonferroni correction for multiple comparisons are indicated on the plot. * p<0.05, **p<0.01. BL: baseline, SWA: slow-wave activity, ZT: zeitgeber time.

**Figure S3.**
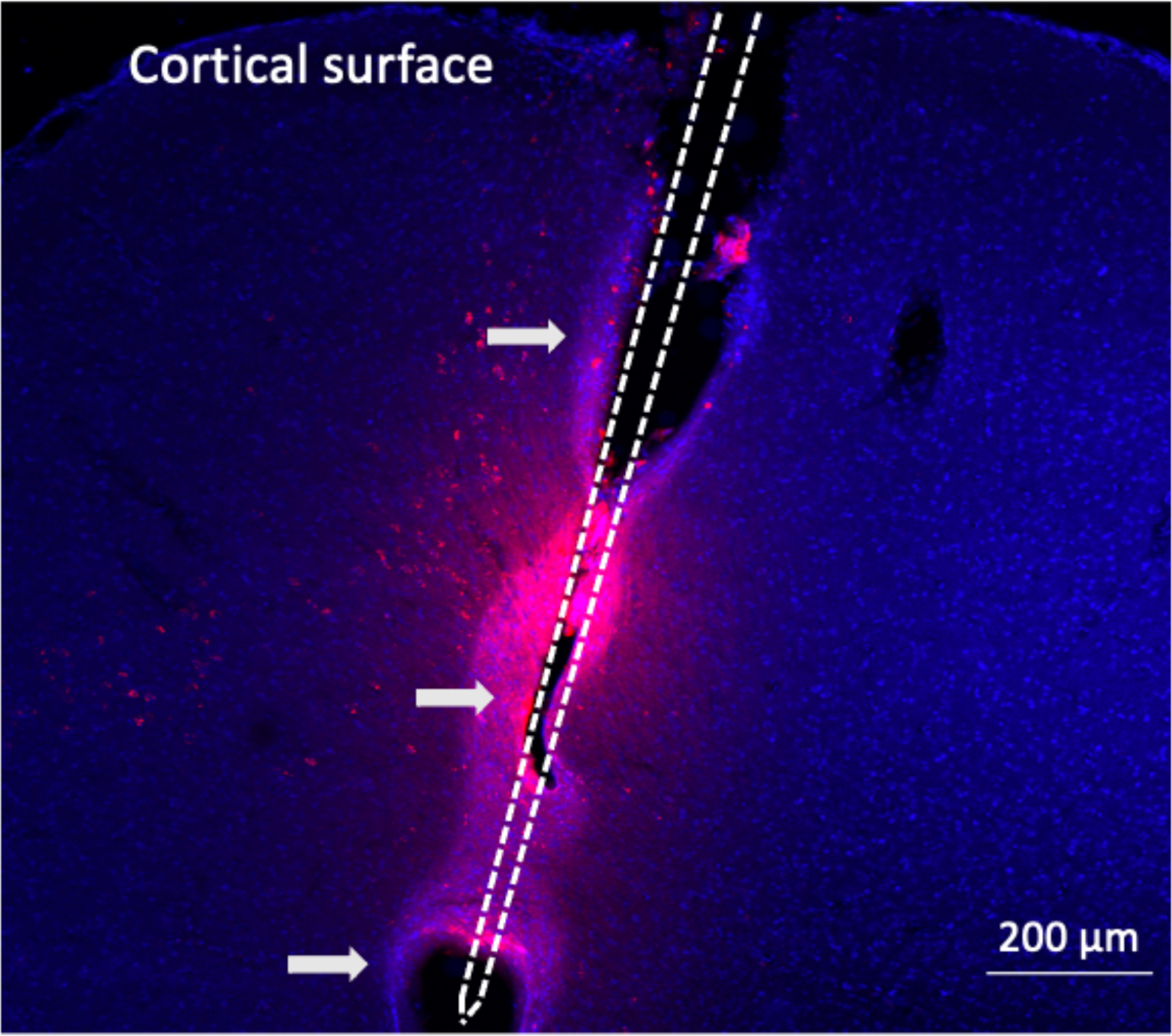
Confocal image of laminar probe placement from one mouse. Grey arrows indicate micro-lesioning sites (see Methods). DiI reported in pink. Cell nuclei in blue (NucBlue).

## Notes

### Competing Interest Statement

The authors have declared no competing interest.

### Summary of Updates

Acknowledgements updated.

